# Comparison of neuromuscular junction dynamics following ischemic and aged skeletal muscle

**DOI:** 10.1101/2021.11.23.469760

**Authors:** Berna Aliya, Mahir Mohiuddin, Jeongmoon J. Choi, Gunjae Jeong, Innie Kang, Hannah Castels, Cade Jones, Young C. Jang

**Author notes:** To whom correspondence should be addressed: Young C. Jang, Ph.D.

## Abstract

Both aging and neuromuscular diseases lead to significant changes in the morphology and functionality of the neuromuscular synapse. Skeletal muscles display a remarkable regenerative capacity, however, are still susceptible to diseases of aging and peripheral nerve perturbations. In this study, we assessed how neuromuscular synapses differ in aged and injured skeletal muscle using an improved neuromuscular junction (NMJ) staining and imaging method. We found that both aged and ischemic skeletal muscle display Wallerian degeneration of the presynaptic motor axons and fragmentation of postsynaptic acetylcholine receptors (AChRs). Quantifiable measurements of various metrics of the NMJs provide a more concrete idea of the dynamics that are occurring in the muscle microenvironment. We questioned whether neuronal degradation precedes myofiber atrophy or vice versa. Previously, it was shown that a cellular crosstalk exists among the motor neurons, myofibers, vasculature, and mitochondria within the muscle microdomain. It is apparent that lack of blood flow to motor neurons in ischemic skeletal muscle disrupts the structure of NMJs, however it is unclear if the aging condition experiences similar dynamics. We demonstrated that both aged and ischemic skeletal muscle demonstrate similar patterns of degeneration, characterized by a smaller percentage overlap of presynaptic and postsynaptic sides, greater fragmentation of AChRs, and a smaller area of AChR clusters. Together, these results reveal high resolution, precise parallels between the aged and ischemic NMJs.

**Impact Statement:** The goal of this study was to assess changes in presynaptic motor neurons and postsynaptic acetylcholine receptors following an ischemic injury model and compare this with an aging model. This was accomplished by characterizing key components of NMJ morphology, including overlap and size of the receptors. There is currently limited research investigating the cellular communication between skeletal muscle fibers and motor neurons. Additionally, there is limited work comparing neuromuscular remodeling in aged and young models. With the substantial prevalence of neuromuscular disorders, especially in the aging population, it is essential to understand nerve-muscle interactions in order to promote increased mobility and improved quality of life in both injury and aging models.

## Introduction

The major functional unit of the neuromuscular system is the motor unit, which consists of the presynaptic motor neuron and all the muscle fibers that it innervates. This unit is essential for producing movement. The synapse at which the alpha motor neuron innervates the postsynaptic acetylcholine receptor (AChR) on the muscle fiber is the neuromuscular junction (NMJ). The role of the NMJ is essential for proper functioning of the entire neuromuscular system and surrounding niche components. This includes, but is not limited to, the vasculature, muscle fibers, muscle satellite cells (MuSCs), and mitochondria.

Blood vessels that are in close proximity to the motor neurons make up the neurovascular system and demonstrate an interdependence of the two tissues to influence one another’s remodeling (Martin & Lewis, 1989). Additionally, ischemia reperfusion injury that disrupts blood flow to skeletal muscles results in deficits in muscle function (Vignaud et al., 2010). This atrophy is further exacerbated by degeneration of the motor axon and the extent of denervation is especially pronounced in aged skeletal muscle (Rowan et al., 2012).

The neuromuscular system is disrupted following disease states as well as with aging. The purpose of this study is to investigate how NMJ dynamics are altered in one particular disease state, critical limb ischemia (CLI), in comparison to uninjured aged muscle. CLI is considered the most severe form of peripheral artery disease (PAD), a condition that affects 8-12 million people in the United States alone and characterized as an atherosclerotic syndrome that causes an obstruction of the peripheral arteries in the legs (Hirsch et al., 2001). Common risk factors for PAD include smoking, diabetes, and age (Hirsch et al., 2001). Although only approximately 1% of the entire PAD patient population is affected by CLI, these patients have the greatest mortality rate of approximately 70% within 10 years of their diagnosis (Varu et al., 2010). With no cure for this condition, current treatments attempt to relieve pain, increase mobility, and reduce cardiovascular risk factors (Slovut & Sullivan, 2008). Therapeutic angiogenic treatments are commonly used in patients to promote new blood vessel formation. Basic fibroblast growth factor is one example of an angiogenic factor that can be injected intramuscularly and results in improved transcutaneous oxygen pressure and ankle brachial index, as well as in the distance walked (Marui et al., 2007). Vascular endothelial growth factor is another angiogenic treatment that has demonstrated surges in collateral vascularization to muscle tissues (Cooke & Losordo, 2015). However, all these therapeutics have had very limited success in promoting a functional neurovascular and neuromuscular system, likely due to the ischemia-induced deterioration of muscle that hosts both blood vessels and motor neurons.

Neuromuscular diseases have demonstrated significant disruptions in many components of the skeletal muscle tissue. Current treatments are insufficient in treating the entire neuromuscular environment and fail to promote a fully functional system. Skeletal muscle exhibits a strong regenerative capacity that may offer a more effective target for regenerative therapeutics. Muscle satellite cells (MuSCs) are stem cells that are essential for the regeneration of skeletal muscle. These cells proliferate and differentiate to fuse into myotubes and finally myofibers. Studies have demonstrated crosstalk evident between MuSCs and motor neurons. In fact, depletion of MuSCs induces a disruption of the morphology of NMJs, and regeneration of NMJs generates an increase in activity of neighboring MuSCs (Liu et al., 2015). Choi et al. (2020) also demonstrated that the crosstalk between MuSCs and motor neurons is diminished following disruption of NMJs. Overall, it is apparent that communication amongst MuSCs and neuronal cells is essential for the maintenance of a functional neuromuscular environment.

CLI and a majority of other neuromuscular disorders present themselves in old age. Thus, it is important to understand the naturally occurring NMJ dynamics that take place in aged skeletal muscle, in order to interpret how an injury model may extrapolate these conditions. It is interesting to note that muscles innervated by brainstem-derived motor neurons are more resistant to the effects of aging (Valdez et al., 2012), in contrast to the spinal nerve innervated skeletal muscles we investigated in this study. NMJ dynamics that occur in aging muscle can be attributed to either disturbances that occur in motor neurons or due to damage in muscle fibers. Aged NMJs have been shown to be relatively stable until experiencing some form of muscle atrophy and regeneration, as indicated by centrally nucleated myofibers (Li et al., 2011). Additionally, the depletion of MuSCs is associated with NMJ deterioration, but not necessarily denervation of the motor neuron (Liu, Klose, et al., 2017). This further supports the hypothesis that some form of paracrine signaling is present between myofiber and motor neurons. However, the particular signaling factor or mechanism that is disrupted in myofibers and subsequently leads to a degeneration of NMJs is not well understood.

There has been limited work investigating the crosstalk between skeletal muscle fibers and motor neurons in both in an ischemic and aged muscle model. Both natural aging and age-induced ischemia demonstrate impairments in myogenesis and neuronal function. Considering the high prevalence of CLI in the aged population, this comparison of the morphological changes in NMJs that occur in ischemic and aged muscle is an important aspect to investigate, although a majority of research conducted for ischemic injury has taken place in young models. By understanding the underlying dynamics that occur in healthy aged muscle, it is easier to understand how these dynamics are altered in an age-induced injury state.

For this study, we utilized Thy1-YFP transgenic mice, a motor neuron specific reporter, for clear visualization of motor neurons in skeletal muscle. Thy1-YFP mice were previously described (Feng et al., 2000). Due to the three-dimensional structure of NMJs, it is difficult to obtain clear images that portray complete dynamics of the synapse. To combat this issue, we obtained longitudinal sections of the NMJs by performing a whole-mount immunofluorescence staining procedure and imaging z-stacks with confocal microscopy. The question we aimed to address is, how do morphological dynamics of presynaptic motor neurons, postsynaptic AChRs, and the synapse as a whole differ between normal aging and the ischemic condition.

Using an improved method of staining combined with microscopy, we are better able to visualize dynamic processes of NMJs in regenerating muscle following ischemic injury. This model can also be applied to more traumatic muscle injury models such as volumetric muscle loss (VML). Comparison of synapse morphology in the aged condition could help with the identification of chemical or molecular factors that may resist age-related changes to NMJs. This can be extended to also encompass neurodegenerative diseases.

We report similar degrees of degeneration and fragmentation of NMJs in aged and ischemic skeletal muscle, indicating parallels between the two conditions. This provides a basis for beginning to understand what signaling pathways and molecular factors are impacted and can be attenuated in aging and neurodegenerative diseases to reduce synaptic damage.

## Methods and materials

### Animals

B6.Cg-Tg(Thy1-YFP)16Jrs/J (#003709) mice purchased from Jackson Laboratories were used for all experiments. Both male and female mice aged 3 to 6 months, considered young adults, were used in a randomized manner all experiments in this study. Aged mice were those 19 months and older. All mice were used according to the protocols approved by the Georgia Institute of Technology Institutional Animal Care and Use Committee.

### Ischemic injury

A murine hindlimb ischemia surgical model was applied to observe the effects of CLI. To perform this, the femoral artery was exposed by making a small incision from the ankle to approximately above the hip. Ligation of the femoral artery was performed using 5-0 sutures between the epigastric and profunda femoris artery. Upon ligating proximal to the tibial artery branching, the artery segment between these ligations was removed, being careful to not sever the femoral nerve. The skin was finally closed using sutures and wound clips. A sham surgery, whereby a similar methodology was used but no ligation, was applied on the contralateral leg. Mice were maintained for 3-56 days following the CLI surgery in individual cages before euthanization by inhalation of CO_2_.

### Tissue histochemistry and immunostaining

Following euthanization of animals, hindlimbs were harvested and fixed in 4% paraformaldehyde for 2 hours at 4°C. The tibialis anterior (TA) and extensor digitorum longus (EDL) muscles were dissected. Prior to staining with antibodies, tissues were blocked for 1 hour using blocking buffer (5% BSA, 0.5% goat serum, 0.5% Triton-X in 1X PBS). Staining media was prepared in blocking buffer using alpha-Bungarotoxin (BTX) conjugated with Alexa Fluor 555 (ThermoFisher, B35451, 1:200 dilution in PBS). Additionally, Alexa Fluor 488-conjugated anti-GFP (ThermoFisher, A21311, 1:250 dilution in PBS) and Alexa Fluor 647-conjugated phalloidin (ThermoFisher, A22287, 1:1000 dilution in PBS) antibodies were added to the secondary staining solution. All secondary antibodies were incubated for 2-3 hours at room temperature. TA samples were separated into bundles and EDL samples were whole mounted using Fluoroshield Mounting Medium with DAPI (Abcam, ab104139) and stored at 4°C.

### Microscope imaging

Samples were imaged on the Zeiss 700 Laser Scanning Confocal Microscope at 63x magnification with oil immersion. Super-resolution images were taken on the Zeiss 780/Elyra PS.1 Superresolution Microscope using Structured Illumination Microscopy (SIM). SIM technique allows for a 115-120 nm resolution in the XY plane and 300 nm in the Z-axis (Badawi & Nishimune, 2020). Z-stack images were taken using both imaging techniques and compiled with Zen Black software (Zeiss).

### Quantification of NMJs

Following microscopic imaging, NMJs were quantified on Fiji/ImageJ using the methodology described by Minty et al. (2020) and the ImageJ Binary Connectivity plugin available at https://blog.bham.ac.uk/intellimic/g-landini-software/ (Landini 2008). Images were cleaned up of any noise and thresholded. Presynaptic nerve terminal perimeter and area measurements were taken, and the complexity was computed. Postsynaptic AChR perimeter and area, endplate diameter, and endplate perimeter and area were taken. From these measurements, compactness, area of synaptic contact, overlap, average area of AChR clusters, and fragmentation were computed. Data was grouped as control, injury time points, or aged. Approximately 15 NMJs were analyzed per sample size from n=3 animals for each group.

### 3D reconstruction of images

Using CZI files obtained from z-stack images taken on confocal microscopy, 3D reconstruction of both NMJs and MuSC was accomplished using Volocity software from Quorom Technologies.

### Statistical Analyses

Statistical analyses in this study were performed in GraphPad Prism 7 software and data is represented as mean +-standard deviation for all figures. A two-tailed unpaired t-test with Welch’s correction was used to compare control and injury groups (Fig. 2) and between control and aged groups (Fig. 3). A p-value of less than 0.05 was considered statistically significant.

## Experimental Results

### Whole mount immunostaining for NMJ exposure

In this study, we utilize an improved method staining and microscopy to better visualize the dynamic processes present in NMJs. By utilizing Thy1-YFP mouse lines combined with alpha-BTX fluorescent tagging, we are able to obtain a more complete and detailed image of both presynaptic and postsynaptic sides of the neuromuscular synapse. Here, we use the EDL muscle that is easily separable into individual bundles and whole mounted. We apply a whole mount immunostaining protocol (Fig. 1A) to hindlimb skeletal muscles. Healthy, young NMJs display significant overlap between motor neurons and post-synaptic AChRs when imaged on laser confocal microscopy (Fig. 1B, top). 3D reconstructions of NMJs reveal a dense concentration of mitochondria (DAPI) at the synapse (Fig. 1B, bottom right). To visualize the postsynaptic AChRs at a higher resolution, super-resolution SIM microscopy was used (Fig. 1B, bottom left). These images reveal the striated pattern present in AChRs, whose nanoscale organization has previously been investigated (York & Zheng, 2017).

**Figure 1:**
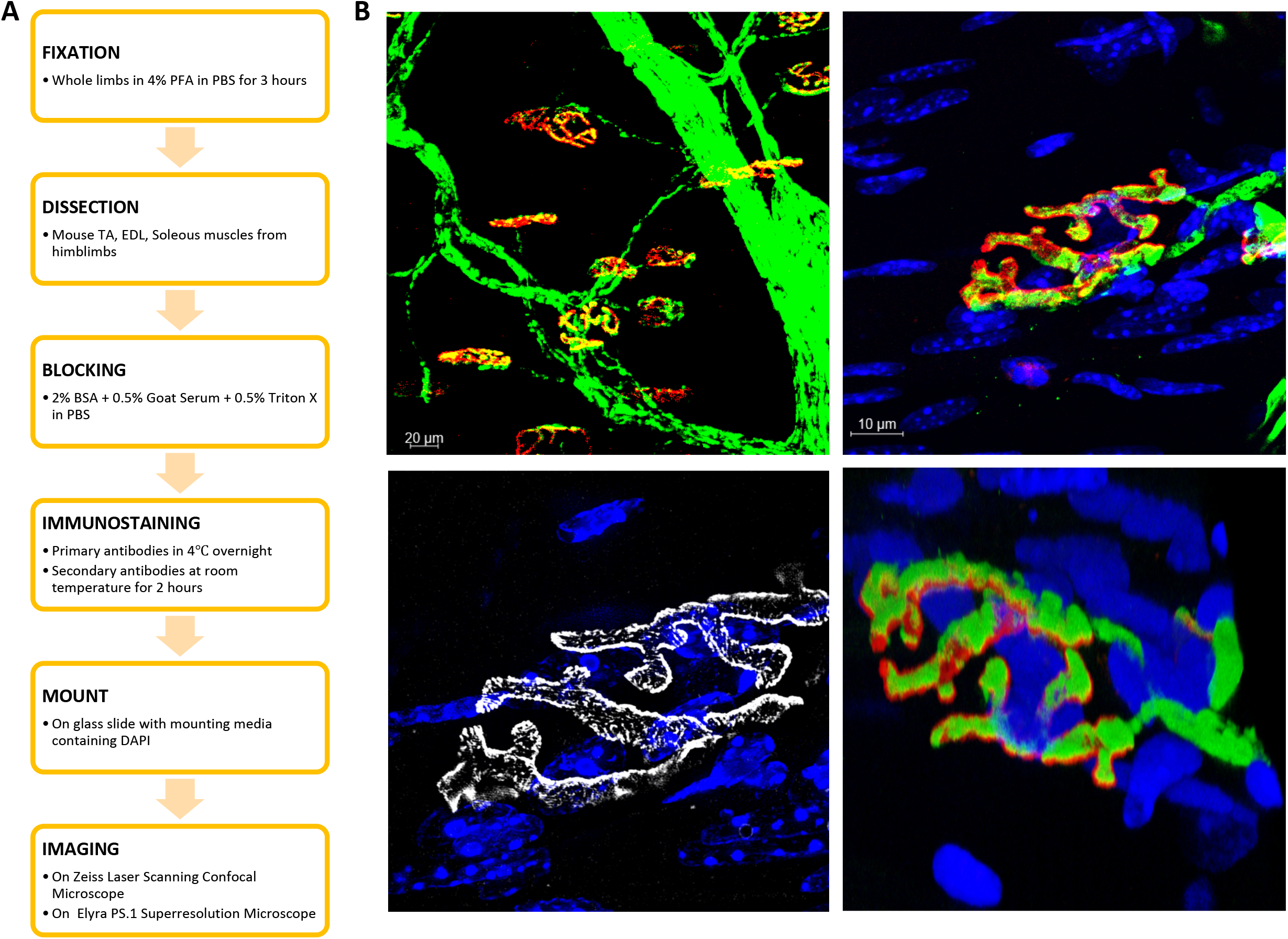
Immunostaining of hindlimb muscles for NMJ detection. **A,** Flowchart demonstrates the protocol used to obtain consistent immunostaining of skeletal muscle fibers. **B**, Immunostaining of young control muscle fibers. Top left image shows a low magnification image of several NMJs distributed along the muscle tissue. antiGFP-488 is shown in green to visualize motor neurons, BTX-555 is shown in red to visualize postsynaptic AChRs. Top right image shows a higher magnification of a single NMJ. Bottom left image shows a super-resolution SIM image of the AChR (white) colabeled with DAPI (blue), visualizing the mitochondria, to demonstrate a nanoscale resolution of the receptor. Bottom right image shows a 3D reconstruction of the same NMJ to provide a more complete image of the cellular environment.

**Figure 2:**
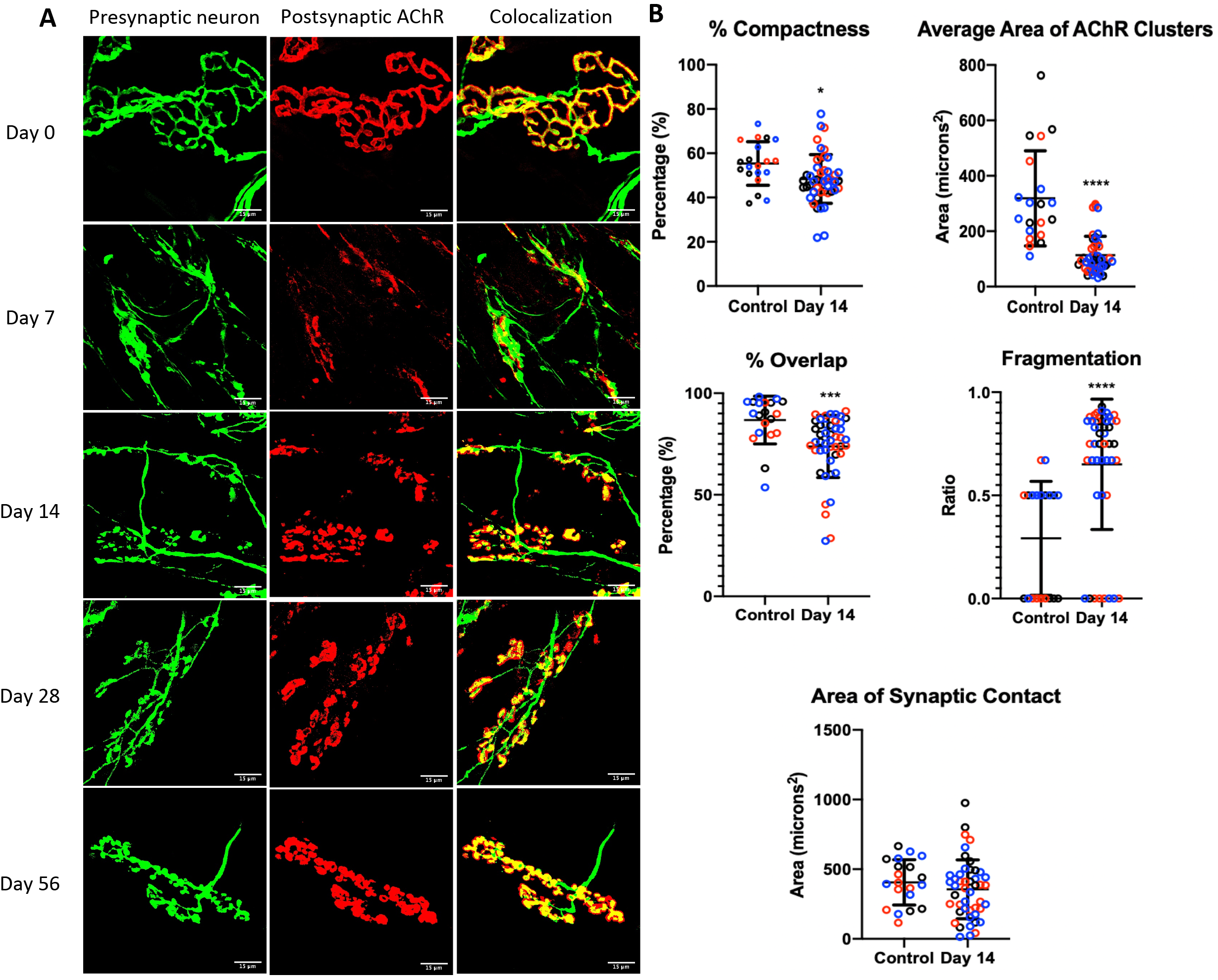
Remodeling of NMJs following ischemic injury. **A,** Confocal microscopy images of NMJs throughout timepoints up to day 56 following ischemia. The greatest fragmentation of AChR and most denervation occurs at day 7 post ischemic injury, followed by a remodeling of the NMJ up to 56 days following injury. **B**, Quantification of NMJs at 14 days post ischemic injury shows significantly less percent compactness, average area of AChR clusters, percent overlap, and significantly greater fragmentation when compared to control. Two tailed unpaired t-test with Welch’s correction: *p = 0.0135, ***p = 0.0005, ****p <0.0001 (n=3). Error bars represent the standard deviation.

**Figure 3:**
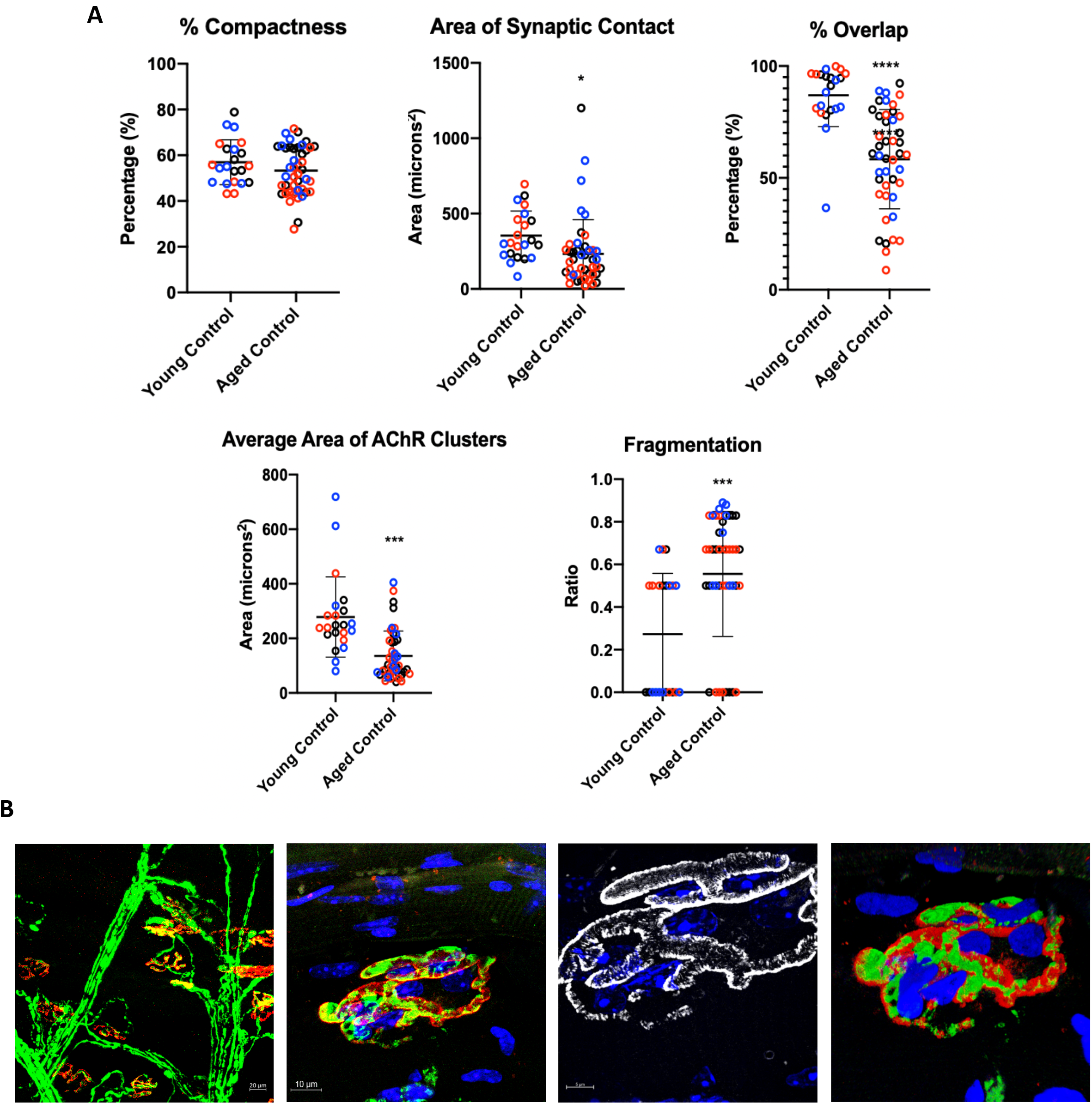
Alterations of neuromuscular junction with aging. **A**, Quantification of aged NMJs shows significantly less area of synaptic contact, percent overlap, average area of AChR clusters, and significantly greater fragmentation when compared to young control. Two tailed unpaired t-test with Welch’s correction: *p = 0.0163, ***p = 0.0005, ****p <0.0001 (n=3). Error bars represent the standard deviation. **B**, Immunostaining of aged control muscle fibers. First image shows a low magnification of several NMJs distributed along the muscle tissue. antiGFP-488 is shown in green to visualize motor neurons, BTX-555 is shown in red to visualize postsynaptic AChRs. Second image shows a higher magnification of a single NMJ. Third image shows a super-resolution SIM image of the AChR (white) colabeled with DAPI (blue), visualizing the mitochondria, to demonstrate a nanoscale resolution of the receptor. Final image shows a 3D reconstruction of the same NMJ to provide a more complete image of the cellular environment.

### NMJ dynamics following ischemic injury

We assessed the morphological changes to the NMJs in Thy1-YFP mice and characterized how the AChR and motor neuron transform following various timepoints post-ischemic injury. The disruption of blood flow induced by ischemic conditions, impacts the neuromuscular system by resulting in Wallerian degeneration of the presynaptic motor neuron and fragmentation of the postsynaptic acetylcholine receptor (Fig. 2A). It appears that the greatest postsynaptic fragmentation and most denervation occurs at day 7 post ischemic injury, followed by a dynamic remodeling of the NMJ up to 56 days following injury (Fig. 2A). Quantitative analysis of the NMJs was performed using the ImageJ/Fiji protocol as described by Minty et al. (2020). We chose to compare only day 14 NMJs to control samples because they demonstrate the most severe phenotype following ischemia, but are still relatively easy to assess, unlike the day 7 samples which reveal too much fragmentation. ImageJ analysis quantified five categories: (1) % compactness, (2) average area of AChR clusters, (3) % overlap, (4) fragmentation ratio indicated by 1.0 being completely fragmented and 0.0 being completely intact, and (5) area of synaptic contact. Approximately 15 NMJs were assessed per biological sample (n=3).

In day 14 ischemic samples, there was overall less percentage compactness of the postsynaptic AChR (48.37%) compared to the control group (55.40%) (Fig. 2B, top left). The average area of postsynaptic clusters is significantly less in the ischemic group, 113.6 microns^2^ (Fig. 2B, top right) and also displays a significantly greater ratio of fragmentation, 0.6504 (Fig. 2B, middle right). Conversely, control NMJs reveal an average area of 318.7 microns^2^ (Fig. 2B, top right), and a fragmentation ratio of 0.2920 (Fig. 2B, middle right). Finally, the control group demonstrates significantly higher percentage overlap of the presynaptic motor neurons and postsynaptic sides with 86.79% overlapped (Fig. 2B, middle left). In contrast, ischemic samples reveal 73.81% overlap (Fig. 2B, middle left). There is no significant difference shown for area of synaptic contact (Fig. 2B, bottom).

### Aged skeletal muscle present chronic NMJ degeneration

Due to the increased likelihood of neurodegenerative diseases, such as CLI, in the aging population, it is important to study the impacts of age on NMJ regeneration. However, studies on NMJ dynamics in aged muscle have been limited. To investigate this, we used immunostaining to visualize NMJs in both aged and young skeletal muscle. Aged control samples display less proper innervation, less overlap of presynaptic and postsynaptic sides, and more Wallerian degeneration (Fig. 3B) when compared to young control samples (Fig. 1B). This is likely due to naturally occurring cycles of denervation and innervation in aged myofibers. Aged samples display a smaller area of synaptic contact, 233.4 microns^2^, versus 354.2 microns^2^ in young samples (Fig. 3B, top middle). There is significantly less percentage overlap of pre- and postsynaptic sides in the aged NMJs, which display 58.37% overlap, in comparison to the young NMJs with 86.96% overlap (Fig. 3B, top right). The average area of AChR clusters in aged NMJs are approximately 135.6 microns^2^ compared to 278.2 microns^2^ in young samples (Fig. 3B, bottom left). Finally, aged samples show a significantly higher ratio of fragmentation, 0.5555, versus 0.2732 in young NMJs (Fig. 3B, bottom right). The percentage compactness of NMJs shows no significant difference between young and aged.

## Discussion

The goal of this study was to characterize the neuromuscular dynamics that occur in ischemic skeletal muscle and compare these findings to the naturally occurring dynamics that occur in aged myofibers. Ischemia-induced disruption of vasculature prevents oxygen and nutrients from being supplied to motor neurons, thereby interrupting proper innervation patterns. A constant supply of oxygen and nutrients is needed for motor neurons to meet the high energy demands of maintaining the membrane potential. Our findings in this study indicate significant changes that occur to the morphology of NMJs following an ischemic injury. We observe Wallerian degeneration of presynaptic motor neurons and fragmentation of the postsynaptic AChRs. Ischemic injury impacts multiple characteristics of NMJ dynamics, including percent compactness, area of AChR clusters, percentage overlap of the synapse, and ratio of fragmentation (Fig. 2). Additional studies that have investigated the role of MuSCs in NMJ regeneration observed an increase in subsynaptic nuclei during the timepoints of greatest denervation following ischemic injury (Schultz, 1978). These nuclei form when MuSCs fuse into myofibers as new nuclei and are important for the formation of AChRs.

Ischemia induced disruption of vasculature induces myofiber atrophy and subsequently damages motor neurons (Mohiuddin et al., 2019). Additionally, it has been shown that capillary-to-fiber ratio increases upon inducing skeletal muscle contraction to stimulate NMJs (Shiragaki-Ogitani et al., 2019). Thus, further supporting the notion of cellular crosstalk evident amongst cellular components of the muscle microenvironment. In both ischemic and aged skeletal muscle, many components of the neuromuscular system are damaged, thereby hindering proper remodeling of cellular domains.

In contrast, aging skeletal muscle goes through repeating cycles of denervation and reinnervation (Aare et al., 2016) leading to destabilized NMJs and chronic muscle atrophy. We observe significant differences in area of synaptic contact, percentage overlap, area of AChR clusters, and fragmentation ratio in aged NMJs (Fig. 3). Scanning electron microscope studies also reveal partially segmented endplates and increased branching of the nerve terminal in the aged NMJs (Fahim et al., 1983). Likely due to an underlying disruption in the cellular crosstalk amongst neuromuscular niche components, proper NMJ formation is hindered. However, the presence of abnormal NMJs in uninjured aged muscle is not unexpected, as it has been shown that NMJ remodeling occurs before any myofiber damage is even present (Deschenes et al., 2010).

The high prevalence of evidence for cellular crosstalk indicates that paracrine signaling likely plays a significant role in NMJ remodeling. One cellular domain that shows potential for understanding the molecular mechanisms responsible for NMJ dynamics is the mitochondria. Previous studies demonstrated that denervation in mouse skeletal muscle results in decreased mitochondrial respiratory capacity and that mitochondrial dysfunction is present prior to any myofiber atrophy (Spendiff et al., 2016). Mitochondria play a significant role in regulating reactive oxygen species (ROS), reactive chemicals that are associated with cellular damage. ROS have shown to be increased in aged muscle mitochondria along with increased myofiber atrophy (Muller et al., 2007). Additionally, we observed that ROS regulatory enzymes, superoxide dismutase 1 and 2 (SOD1 and SOD2) are disrupted in ischemic conditions, as indicated by a significant decline in specific activity of SOD2, a measure of the catalytic activity of the enzyme per total quantity (Fig. S1). This decline in catalytic activity correlates with the NMJ regeneration patterns present from day 7 to day 28 (Fig. 2), where we observe the presynaptic and postsynaptic sides attempt to reinnervate as SOD2 enzyme content increases but activity decreases. This alludes to the presence of other factors in the muscle microenvironment that are likely hindering proper functionality of ROS regulatory enzymes. Stimulation of NMJs was shown to increase the number of oxidative myofibers, indicating that injured limbs improve their oxygen usage (Shiragaki-Ogitani et al., 2019). It is also evident that mitochondria are essential for the remodeling of NMJs due to their high concentration on both presynaptic and postsynaptic sides of the muscle synapse (Rygiel et al., 2016). Substantial changes in mitochondria morphology and protein composition are evident in aged rats, characterized by swelling and mitochondrial fusion (García et al., 2013).

Likewise, mitochondrial dysfunction has been associated with age-related muscle atrophy, or sarcopenia. Primarily, mitochondrial energy metabolism pathways, such as oxidative phosphorylation and the Krebs cycle, have the most genes downregulated when assessing global gene expression profiles (Ibebunjo et al., 2013). For our future studies, it would be interesting to investigate the content and activity of these proteins in young and aged models and compare them to the ischemic condition. This would allow for a more comprehensive understanding of what molecular mechanisms contribute to age-related neuromuscular changes. MuSC-stemmed mitochondria show promising results in improving bioenergetic function (Mohiuddin et al., 2020), thus are a promising therapeutic target for neuromuscular conditions that exacerbate mitochondrial dysfunction.

## Conclusion

In summary, we demonstrate the prevalence of NMJ dynamics following ischemic and aged skeletal muscle. Cellular crosstalk amongst the vascular system, motor neurons, muscle fibers, MuSCs, and mitochondria supports the notion of paracrine signaling and the likely influences that biochemical changes in one cell will have on a neighboring cell (Fig. 4). By understanding the underlying NMJ dynamics in aged muscle and the biochemical changes that cause this, we can better hypothesize the molecular mechanisms that are responsible for neuromuscular degeneration in age-induced diseases, such as CLI.

**Figure 4:**
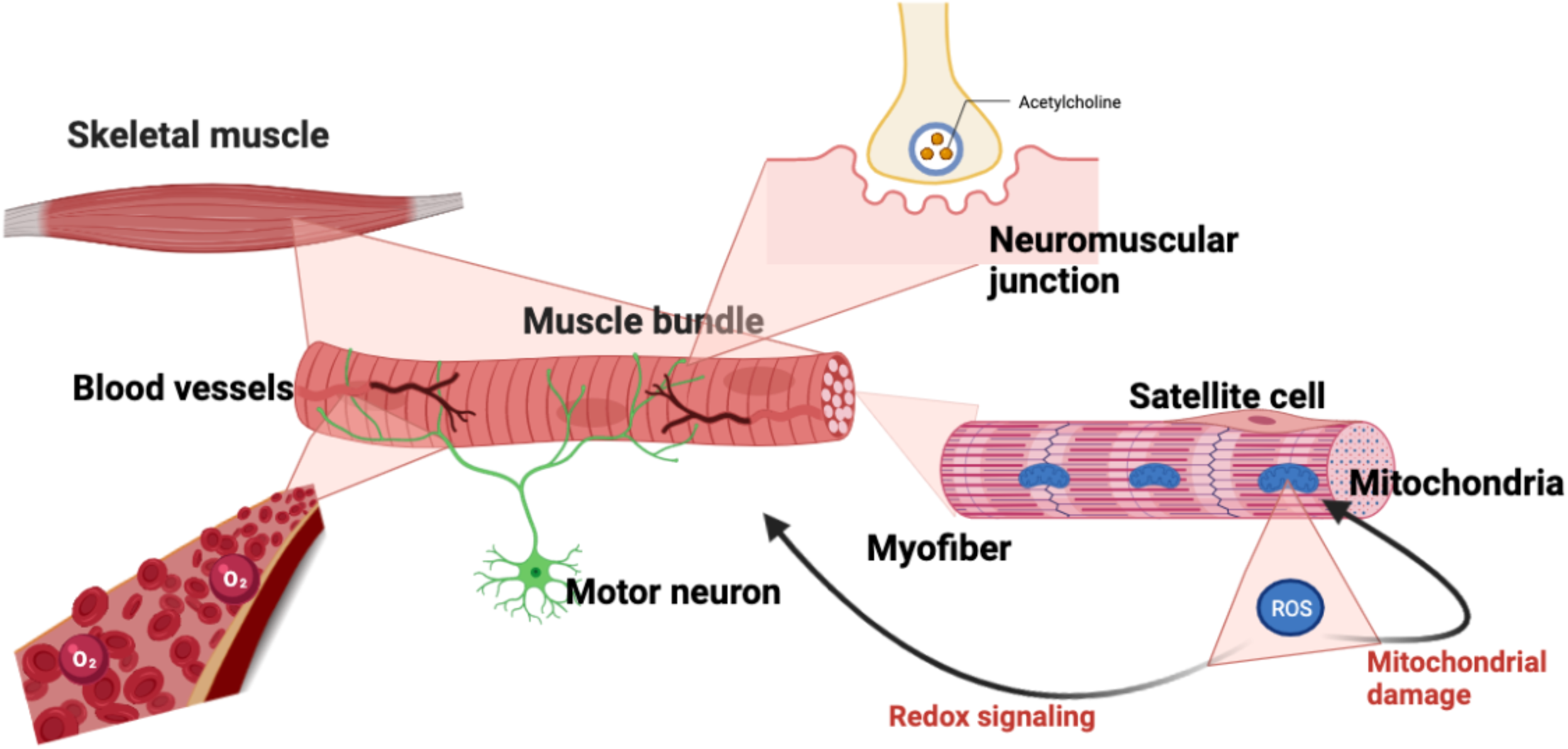
Crosstalk amongst cells of muscle microenvironment. Infographic representing various cellular components present in the muscle niche. This encompasses the vascular system, motor neurons, muscle fibers, muscle stem cells, and mitochondria. Created with BioRender.com.

## Acknowledgements

We thank the Physiological Research Laboratory and the core facilities at the Parker H. Petit Institute for Bioengineering and Bioscience. This work was supported by the National Institutes of Health under award numbers R21AR072287 (Y.C.J.), Department of Defense W81XWH-20-1-0336 (Y.C.J. and C.A.A.), and S&R Foundation (Y.C.J.). The content is solely the responsibility of the authors and does not necessarily represent the official views of the National Institutes of Health.

## References

1. Aare, S., Spendiff, S., Vuda, M., Elkrief, D., Perez, A., Wu, Q., Mayaki, D., Hussain, S. N. A., Hettwer, S., & Hepple, R. T. (2016). Failed reinnervation in aging skeletal muscle. Skeletal Muscle, 6(1). https://doi.org/10.1186/s13395-016-0101-y

2. Badawi, Y., & Nishimune, H. (2020). Super-resolution microscopy for analyzing neuromuscular junctions and synapses. Neuroscience Letters, 715. https://doi.org/10.1016/j.neulet.2019.134644

3. Choi, J. J., Shin, E. J., Han, W. M., Anderson, S. E., Mohiuddin, M., Lee, N. H., Tran, T., Nakhai, S., Jeong, H., Shcherbina, A., Jeong, G., Oh, D. G., Weinstock, L. D., Sankar, S. B., Ogle, M. E., Katsimpardi, L., Rao, T. N., Wood, L., Aguilar, C. A.,… Jang, Y. C. (2020). Regenerating motor neurons prime muscle stem cells for myogenesis by enhancing protein synthesis and mitochondrial bioenergetics. BioRxiv, 2020.05.24.113456. https://doi.org/10.1101/2020.05.24.113456

4. Cooke, J. P., & Losordo, D. W. (2015). Modulating the Vascular Response to Limb Ischemia. Circulation Research, 116(9). https://doi.org/10.1161/CIRCRESAHA.115.303565

5. Deschenes, M. R., Roby, M. A., Eason, M. K., & Harris, M. B. (2010). Remodeling of the neuromuscular junction precedes sarcopenia related alterations in myofibers. Experimental Gerontology, 45(5). https://doi.org/10.1016/j.exger.2010.03.007

6. Fahim, M. A., Holley, J. A., & Robbins, N. (1983). Scanning and light microscopic study of age changes at a neuromuscular junction in the mouse. Journal of Neurocytology, 12(1). https://doi.org/10.1007/BF01148085

7. Feng, G., Mellor, R. H., Bernstein, M., Keller-Peck, C., Nguyen, Q. T., Wallace, M., Nerbonne, J. M., Lichtman, J. W., & Sanes, J. R. (2000). Imaging Neuronal Subsets in Transgenic Mice Expressing Multiple Spectral Variants of GFP. Neuron, 28(1). https://doi.org/10.1016/S0896-6273(00)00084-2

8. García, M. L., Fernández, A., & Solas, M. T. (2013). Mitochondria, motor neurons and aging. Journal of the Neurological Sciences, 330(1), 18–26. https://doi.org/10.1016/J.JNS.2013.03.019

9. Hirsch, A. T., Criqui, M. H., Treat-Jacobson, D., Regensteiner, J. G., Creager, M. A., Olin, J. W., Krook, S. H., Hunninghake, D. B., Comerota, A. J., Walsch, M. E., McDermott, M. M., & Hiatt, W. R. (2001). Peripheral Arterial Disease Detection, Awareness, and Treatment in Primary Care. JAMA, 286(11). https://doi.org/10.1001/jama.286.11.1317

10. Ibebunjo, C., Chick, J. M., Kendall, T., Eash, J. K., Li, C., Zhang, Y., Vickers, C., Wu, Z., Clarke, B. A., Shi, J., Cruz, J., Fournier, B., Brachat, S., Gutzwiller, S., Ma, Q., Markovits, J., Broome, M., Steinkrauss, M., Skuba, E.,… Glass, D. J. (2013). Genomic and Proteomic Profiling Reveals Reduced Mitochondrial Function and Disruption of the Neuromuscular Junction Driving Rat Sarcopenia. Molecular and Cellular Biology, 33(2). https://doi.org/10.1128/MCB.01036-12

11. Landini G. Advanced shape analysis with ImageJ. Proceedings of the Second ImageJ User and Developer Conference, Luxembourg, 6-7 Nov, 2008. p116–121. ISBN 2-919941-06-2. Plugins available from https://blog.bham.ac.uk/intellimic/g-landini-software/

12. Li, Y., Lee, Y. i., & Thompson, W. J. (2011). Changes in Aging Mouse Neuromuscular Junctions Are Explained by Degeneration and Regeneration of Muscle Fiber Segments at the Synapse. Journal of Neuroscience, 31(42). https://doi.org/10.1523/JNEUROSCI.3590-11.2011

13. Liu, W., Klose, A., Forman, S., Paris, N. D., Wei-Lapierre, L., Corté S-Lopé Z, M., Tan, A., Flaherty, M., Miura, P., Dirksen, R. T., & Chakkalakal, J. v. (n.d.). Loss of adult skeletal muscle stem cells drives age-related neuromuscular junction degeneration. https://doi.org/10.7554/eLife.26464.001

14. Liu, W., Wei-Lapierre, L., Klose, A., Dirksen, R. T., & Chakkalakal, J. v. (n.d.). Inducible depletion of adult skeletal muscle stem cells impairs the regeneration of neuromuscular junctions. https://doi.org/10.7554/eLife.09221.001

15. Martin, P., & Lewis, J. (1989). Origins of the neurovascular bundle: interactions between developing nerves and blood vessels in embryonic chick skin. The International Journal of Developmental Biology, 33(3).

16. Marui, A., Tabata, Y., Kojima, S., Yamamoto, M., Tambara, K., Nishina, T., Saji, Y., Inui, K., Hashida, T., Yokoyama, S., Onodera, R., Ikeda, T., Fukushima, M., & Komeda, M. (2007). A Novel Approach to Therapeutic Angiogenesis for Patients With Critical Limb Ischemia by Sustained Release of Basic Fibroblast Growth Factor Using Biodegradable Gelatin Hydrogel An Initial Report of the Phase I-IIa Study. Circulation Journal, 71(8). https://doi.org/10.1253/circj.71.1181

17. Minty, G., Hoppen, A., Boehm, I., Alhindi, A., Gibb, L., Potter, E., Wagner, B. C., Miller, J., Skipworth, R. J. E., Gillingwater, T. H., & Jones, R. A. (2020). ANMJ-morph: A simple macro for rapid analysis of neuromuscular junction morphology. Royal Society Open Science, 7(4). https://doi.org/10.1098/rsos.200128

18. Mohiuddin, M., Choi, J. J., Lee, N. H., Jeong, H., Anderson, S. E., Han, W. M., Aliya, B., Peykova, T. Z., Verma, S., García, A. J., Aguilar, C. A., & Jang, Y. C. (2020). Transplantation of Muscle Stem Cell Mitochondria Rejuvenates the Bioenergetic Function of Dystrophic Muscle. BioRxiv, 2020.04.17.017822. https://doi.org/10.1101/2020.04.17.017822

19. Mohiuddin, M., Lee, N. H., Moon, J. Y., Han, W. M., Anderson, S. E., Choi, J. J., Shin, E., Nakhai, S. A., Tran, T., Aliya, B., Kim, D. Y., Gerold, A., Hansen, L. M., Taylor, W. R., & Jang, Y. C. (2019). Critical Limb Ischemia Induces Remodeling of Skeletal Muscle Motor Unit, Myonuclear-, and Mitochondrial-Domains. Scientific Reports, 9(1). https://doi.org/10.1038/s41598-019-45923-4

20. Muller, F. L., Song, W., Jang, Y. C., Liu, Y., Sabia, M., Richardson, A., & van Remmen, H. (2007). Denervation-induced skeletal muscle atrophy is associated with increased mitochondrial ROS production. American Journal of Physiology-Regulatory, Integrative and Comparative Physiology, 293(3). https://doi.org/10.1152/ajpregu.00767.2006

21. Rowan, S. L., Rygiel, K., Purves-Smith, F. M., Solbak, N. M., Turnbull, D. M., & Hepple, R. T. (2012). Denervation Causes Fiber Atrophy and Myosin Heavy Chain Co-Expression in Senescent Skeletal Muscle. PLoS ONE, 7(1). https://doi.org/10.1371/journal.pone.0029082

22. Rygiel, K. A., Picard, M., & Turnbull, D. M. (2016). The ageing neuromuscular system and sarcopenia: a mitochondrial perspective. The Journal of Physiology, 594(16). https://doi.org/10.1113/JP271212

23. Schultz, E. (1978). Changes in the satellite cells of growing muscle following denervation. The Anatomical Record, 190(2). https://doi.org/10.1002/ar.1091900212

24. Shiragaki-Ogitani, M., Kono, K., Nara, F., & Aoyagi, A. (2019). Neuromuscular stimulation ameliorates ischemia-induced walking impairment in the rat claudication model. Journal of Physiological Sciences, 69(6). https://doi.org/10.1007/s12576-019-00701-9

25. Slovut, D. P., & Sullivan, T. M. (2008). Critical limb ischemia: medical and surgical management. Vascular Medicine, 13(3). https://doi.org/10.1177/1358863X08091485

26. Spendiff, S., Vuda, M., Gouspillou, G., Aare, S., Perez, A., Morais, J. A., Jagoe, R. T., Filion, M.-E., Glicksman, R., Kapchinsky, S., MacMillan, N. J., Pion, C. H., Aubertin-Leheudre, M., Hettwer, S., Correa, J. A., Taivassalo, T., & Hepple, R. T. (2016). Denervation drives mitochondrial dysfunction in skeletal muscle of octogenarians. The Journal of Physiology, 594(24). https://doi.org/10.1113/JP272487

27. Varu, V. N., Hogg, M. E., & Kibbe, M. R. (2010). Critical limb ischemia. Journal of Vascular Surgery, 51(1). https://doi.org/10.1016/j.jvs.2009.08.073

28. Vignaud, A., Hourde, C., Medja, F., Agbulut, O., Butler-Browne, G., & Ferry, A. (2010). Impaired Skeletal Muscle Repair after Ischemia-Reperfusion Injury in Mice. Journal of Biomedicine and Biotechnology, 2010. https://doi.org/10.1155/2010/724914

29. York, A. L., & Zheng, J. Q. (2017). Super-Resolution Microscopy Reveals a Nanoscale Organization of Acetylcholine Receptors for Trans-Synaptic Alignment at Neuromuscular Synapses. Eneuro, 4(4). https://doi.org/10.1523/ENEURO.0232-17.2017

