## Supplemental Information for "Comparison of neuromuscular junction dynamics following ischemic and aged skeletal muscle"

**Figure S1.**

**A**

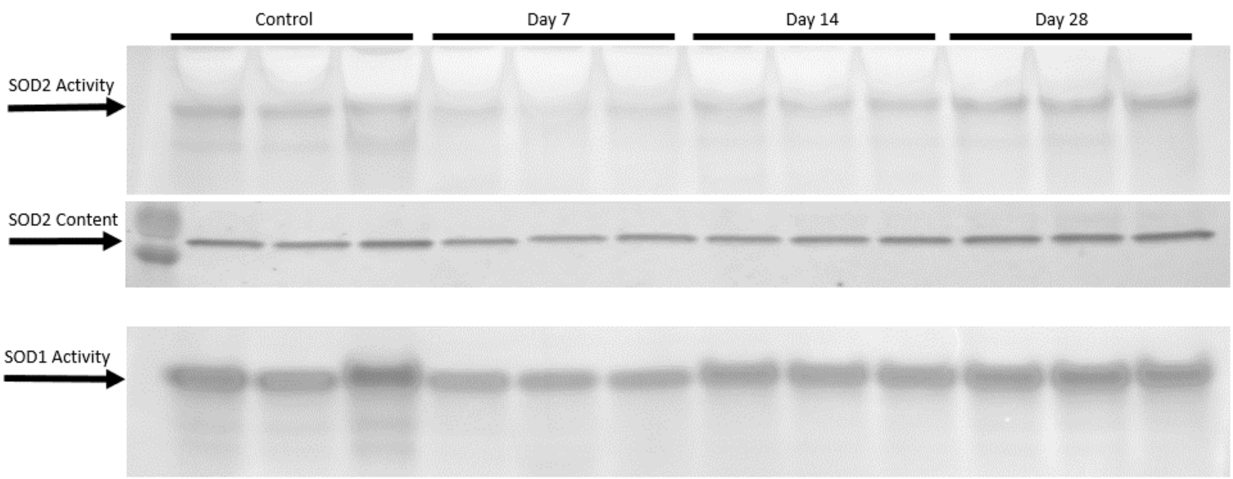


**B**

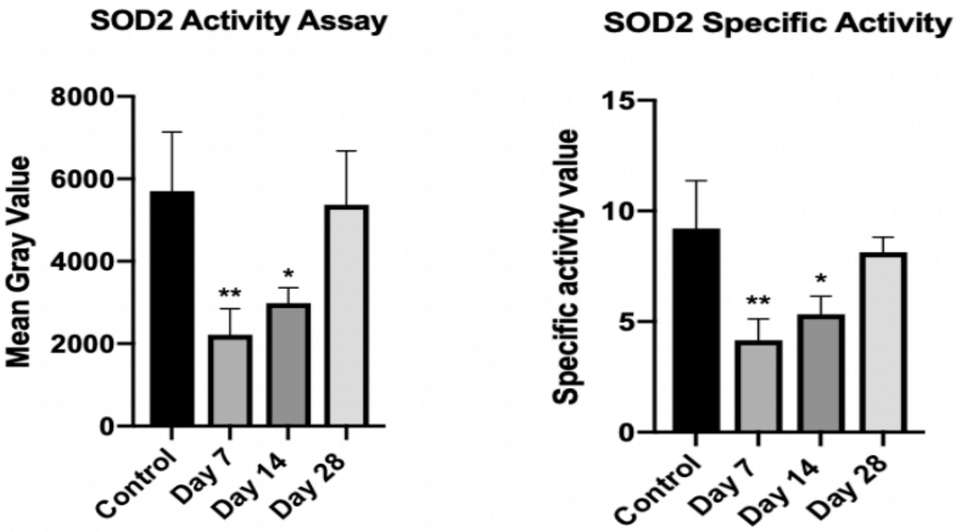


**Supplemental Figure S1: SOD enzyme activity and content analysis in ischemic skeletal muscle. A,** SOD1 and SOD2 activity assays on skeletal muscle homogenates following various time point post-ischemic injury. The greatest decreased in SOD2 activity is evident 7 days post injury. SOD2 western blot indicates a gradual increase in SOD2 content from day 7 to day 28. B, Quantification of SOD2 enzymes demonstrates the greatest decline in SOD2 activity and specific activity at 7 days post ischemic injury. Ordinary one-way ANOVA with Dunnett's multiple comparisons test. On activity assay: *p = 0.0310, **p = 0.0086. On specific activity: *p = 0.0159, **p = 0.0036. Error bars represent the standard deviation.
